# FreeClimber: Automated quantification of climbing performance in *Drosophila*, with examples from mitonuclear genotypes

**DOI:** 10.1101/2020.07.04.187898

**Authors:** Adam N. Spierer, Denise Yoon, Chen-Tseh Zhu, David M. Rand

**Author notes:** **Corresponding authors:** David M. Rand, 80 Waterman St, Providence, RI 02912, Adam N. Spierer, 80 Waterman St., Providence, RI 02912.

## Abstract

Negative geotaxis (climbing) performance is a useful metric for quantifying *Drosophila* health and vigor. Manual methods to quantify climbing performance are slow, tedious, and may be systematically biased, while available computational methods have inflexible hardware or software requirements. We present an alternative: FreeClimber. This open source, Python-based pipeline subtracts a video’s static background to improve spot detection for moving flies in heterogeneous backgrounds. FreeClimber calculates a cohort’s velocity as the slope of the most linear portion of a mean-vertical position vs. time plot. It can run from a graphical user interface for parameter optimization or a command line interface for high-throughput and automated batch processing. It outputs calculated slopes, spot locations for follow up analyses such as tracking, and several visualizations and diagnostic plots. We demonstrate FreeClimber’s utility in a longitudinal study for endurance exercise performance in *Drosophila* using six distinct mitochondrial haplotypes paired with a common *w*^*1118*^ nuclear background.

**Summary statement:** FreeClimber quantifies the climbing velocity for a group of flies, eliminating systematic biases associated with traditional manual methods in a high throughput and automated (graphical and/or command line-based) platform.

## Introduction

The *Drosophila* model system provides a rich set of genetic resources to explore the functional bases of traits at organismal, cellular and molecular levels (Bellen et al., 2011; Chow and Reiter, 2017; Lenz et al., 2013; Mackay et al., 2012). One of the most common *Drosophila* health metrics is locomotor capacity, easily measured using a negative geotaxis (climbing) assay (Gargano et al., 2005; Jones and Grotewiel, 2011). Here, flies are gently knocked to the bottom of a vial and are recorded by video as they instinctively climb upward (Ganetzky and Flanagan, 1978; Gargano et al., 2005). Climbing performance is often reported as some measure of the flies’ position vs. time, such as mean position at a time cutoff (Gargano et al., 2005; Lavoy et al., 2018) or time until a percentage of flies reach a set height (Ma et al., 2014; Podratz et al., 2013; Tsai et al., 2016; Xu et al., 2008).

The climbing assay’s popularity is largely due to its accessibility. Experimental setups are easily engineered from common laboratory items, meaning they are relatively inexpensive to implement. Data collection is straightforward, only requiring simple image capture tools and basic software available on most computers. However, this assay’s simplicity is beset by its tedious and time-consuming nature.

Several publications describe protocols for automating the conversion of visual media into data, but these are not always accessible to the general community. Some of these platforms are detectors, while others are trackers. Detectors identify the x,y-coordinates of spots (flies) across frames, which can be evaluated as a function of position vs. time (e.g. RflyDetection R module (Cao et al., 2017) and an ImageJ-based approach (Podratz et al., 2013)). Trackers build on this with predictive linking to connect spots between frames based on their proximity and likelihood of being connected (e.g. the Hillary Climber tracks single flies in individual vials (Willenbrink et al., 2016), the iFly system tracks multiple flies in a single vial (Kohlhoff et al., 2011), and the DaRT system tracks multiple flies in multiple enclosures (Faville et al., 2015; Taylor and Tuxworth, 2019)). Trackers are challenging to automate because they generally require supervision to discern erratic vertical motions (jumps and falls) or paths that laterally intersect with other flies (collision on the same plane or eclipse on separate planes)) (Chenouard et al., 2014). Additionally, published methods for both detectors and trackers often require a homogeneous background, a custom setup, code from proprietary languages (MATLAB), and/or are only made available locally. Because of these and other factors, no platform is widely accepted by the *Drosophila*-research community, despite the assay’s ubiquity.

We created FreeClimber to address some of these major issues, correct for common biases in traditional manual approaches (such as irregular starting heights), and facilitate the generation of accurate and repeatable data that are more representative of the flies’ motion. This Python 3-based platform can be run interactively, via Graphical User Interface (GUI), or through a command line interface for automated and high-throughput batch processing. FreeClimber utilizes an efficient background subtraction step, so it performs respectably with heterogeneous backgrounds. Additionally, our detector implements a local linear regression for calculating a group’s velocity (Olito et al., 2017), which captures an objective metric of fly climbing that may be challenging with traditional manual analyses. Finally, we demonstrate the utility of our platform for longitudinal *Drosophila* screens analyzing two original data sets of exercise conditioned and unconditioned mitochondrial-nuclear (mito-nuclear) introgression flies. We highlight FreeClimber’s ability to quantify subtle differences in phenotype across longitudinal and sample-rich studies, like those frequently conducted in *Drosophila* research.

## Materials and Methods

### Drosophila husbandry and generation of lines

Six mitochondrial haplotypes (mtDNAs or mitotypes) were derived from three different *Drosophila* species: *D. melanogaster* (subtypes: *OregonR* (OreR) and *Zimbabwe53* (Zim)), *D. simulans* (subtypes: *siI* and *siII*), *D. mauritiana* (subtype: *mall*), and *D. yakuba* (subtype: *yakuba*) (Montooth et al., 2010; Mossman et al., 2016; Zhu et al., 2014). These mitotypes were each placed on a common, *D. melanogaster w*^*1118*^ nuclear background using balancer chromosome crosses and subsequent recurrent male backcrossing using *w*^*1118*^ males (Zhu et al., 2014), with *D. simulans maII* and *D. yakuba* lines created by microinjection of cytoplasm donor into an egg (Ma et al., 2014).

Stocks were density controlled for two generations whereby 20 females and 20 males were allowed to lay eggs for three days per brood. Fly cultures were held at 25°C on standard lab food (Mossman et al., 2016) and maintained on a 12h:12h light:dark schedule. Adult males were collected three days post-eclosion using light CO_2_ anesthetization and separated into vials of 20 flies. Flies were given 24 hours to recover and transferred to new food every day. Testing day number in the longitudinal experiment does not include the three days post-eclosion before exercise training began.

### Video recording set up and recording

While FreeClimber does not require a specialized set up, we used one to standardize our recording environment (Fig. S1) and precisely time video capture (Fig. S2). The main component is a custom polycarbonate climbing rig, comprised of a fixed base with aluminum rails that a mobile chassis could slide along (Fig. S1B). This chassis held six narrow glass vials, evenly spaced, that could be raised and subsequently dropped from a pre-designated height (7 cm) to control for the amount of force applied to all vials when beginning the assay. The base of the rig was mounted on a MakerBeam frame (MakerBeam, Utrecht, Netherlands) and held in place with a setscrew. The frame also held a LED-light board (Huion model L4S, 10.7 lumens/inch^2^; Fuyong, Bao’an District, Shenzhen, China) to backlight the flies. An 8 megapixel PiCamera (V2) attached to a Raspberry Pi 3 (Model B+) (Raspberry Pi Foundation, Cambridge, England, UK) recorded videos from a fixed distance to standardize the video recording parameters.

We also wired a phototrigger to begin video recording as the climbing rig dropped and the assay began. The amount of light emitted from the LED light box was measured by a photoresistor (SEN-09088; Sparkfun Electronics, Niwot, Colorado, USA) and passed through an analog-digital converter (MCP3000; Adafruit Industries, New York, New York, USA). Both were wired to the Raspberry Pi through the General Processing Input Output (GPIO) pins (Fig. S2A). The photoresistor was placed on a frame rail close to the LED light box, separated by an opaque tab attached to climbing rig chassis (Fig. S2B) such that the light path was uninterrupted when the chassis was raised and blocked with it was lowered. After raising and dropping the climbing rig’s chassis, a five second H264 video was recorded at 29 frames per second. Flies were given ten seconds from the end of the video recording to recover before they were tested again. Three technical replicates were recorded for each sex-genotype combination.

### Overview of FreeClimber modes

The platform can be run in two modes: a Graphical User Interface (GUI) for optimizing detection parameters of a single video, and a command line interface for high-throughput batch processing of many videos with a common set of detection parameters (Fig. 1A). Please refer to the GitHub repository for a complete guide on installation and usage, as well as tips and tricks for increasing data quality: https://github.com/adamspierer/FreeClimber/tree/JEB_release. The most current version is available at: https://github.com/adamspierer/FreeClimber/.

**Figure 1.**
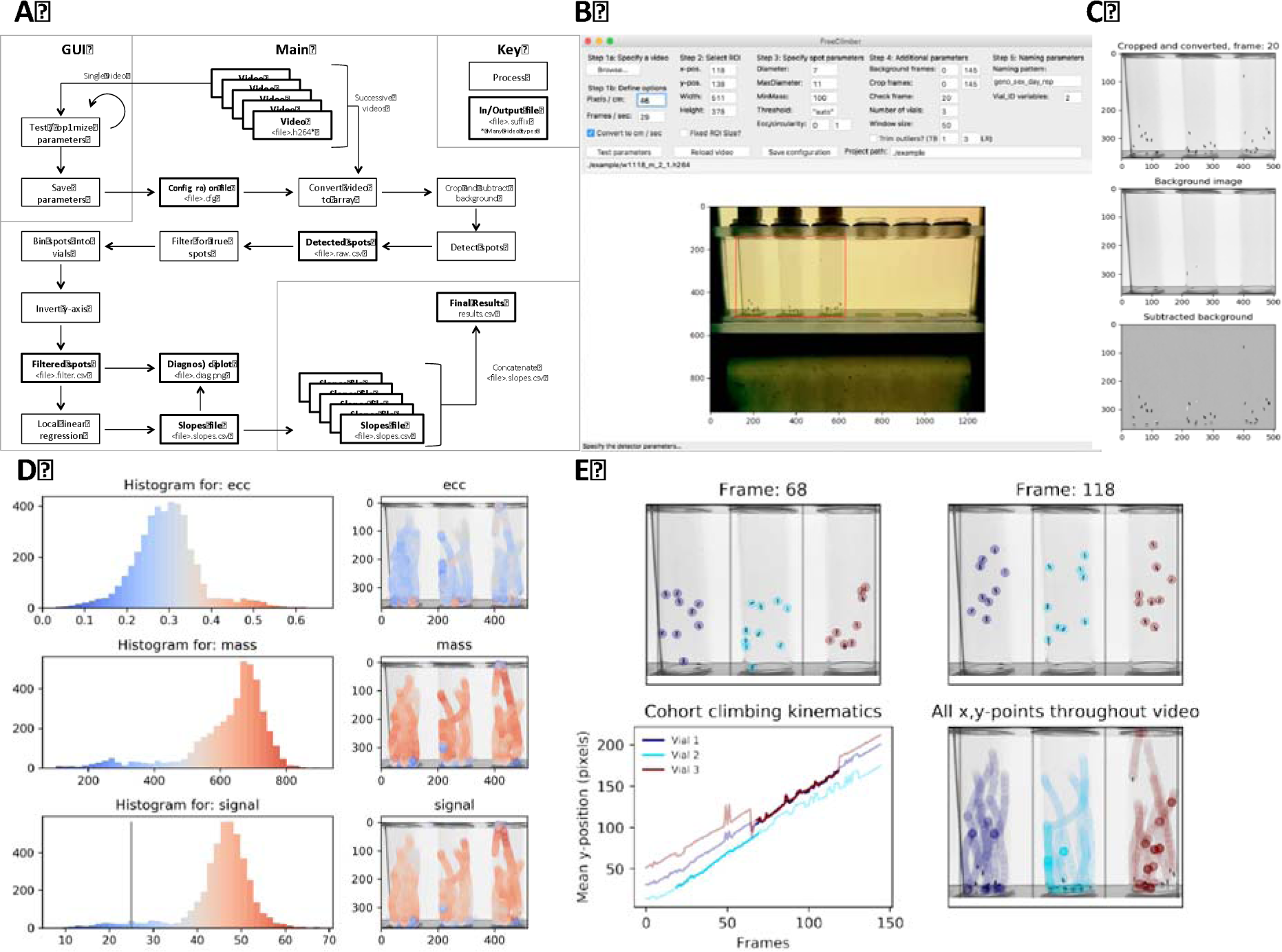
Overview of FreeClimber platform. (A) Flow diagram FreeClimber’s processes. The Graphical User Interface (GUI) is designed for parameter optimization and the command line tool is designed for high throughput processing. (B) Screenshot of GUI with Region of Interest (ROI) outlined in red. (C) Visualization of background subtraction shows the original image recolored and cropped (top), static background matrix (middle), and final subtracted image used for spot detection (bottom). (D) Optimization plots visualize the distribution and location of each spot and its respective metric: ecc (eccentricity, roundness), mass, and signal. (E) Visualization of spot locations in the first and last frames of the most linear segment of all flies’ climbing (top), the most linear portion (darker shade) of the mean-vertical position vs. frame curve plotted over all frames (lighter shade) for an indicated vial (lower-left), and all x,y-coordinates throughout the video (lower-right).

### Video preprocessing and background subtraction

Videos are read into integer-based n-dimensional arrays (*nd*-array) using the FFmpeg-python package (v.4.0.4; https://github.com/kkroening/ffmpeg-python). Following user-defined parameters, videos are cropped for the appropriate frame range and positional region of interest (ROI) (Fig. 1B) before being converted to gray scale. A matrix representing the static background is calculated from the median pixel intensity of each x,y-coordinate across a user-defined number of frames (default is all frames). This background matrix is subtracted from each individual frame’s pixel intensity matrix, resulting in a new *nd*-array corresponding with only regions of movement (flies) in the video (Fig. 1C).

### Detector optimization

The background-subtracted frames are passed to a Python-implementation of the Crocker and Weeks particle-tracking algorithm trackpy (v.0.4.2)(Crocker and Grier, 1996) for spot detection. Spots are identified from clusters of pixels that meet user-defined parameters for the expected spot diameter (diameter) and maximum diameter (maxsize). They must also exceed filtering thresholds for spot roundness (ecc; eccentricity), minimum integrated brightness (minmass), signal threshold (threshold) to be considered a true spot. These parameters can be visualized for all spots in a video to assist in parameter optimization (Fig. 1D). Spots are considered ‘True’ if they pass thresholds for the ecc, minmass, and signal threshold and are visualized in the file with the ‘*spot_check*.*png*’ suffix, created automatically from the GUI or with the ‘-- optimization_plots’ argument in the command line interface. While the signal threshold can be provided by the user, FreeClimber can also calculate an appropriate one using the SciPy (v.1.4.1) functions: peak_prominences and find_peaks. This method looks for the local minimum between two signal peaks in a histogram of spots signal values, or is assigned to be one half of the value of the global maximum if there is only one.

### Spot detection

As the program runs, a data frame containing the spatio-temporal data for each spot and its accompanying metrics is saved with the *’raw*.*csv*’ file suffix. This file can be used as an input for with trackpy to track spots (see Step 3: “Link features into particle trajectories”, http://soft-matter.github.io/trackpy/dev/tutorial/walkthrough.html).

Spot coordinates are transformed and processed for more accurate estimation of group climbing velocity. The raw data set is filtered for only true spots, described above. Y-coordinates are inverted to account for images being indexed from upper-left to lower-right, instead of lower-left to upper-right (Fig. 1E). Vials are assigned by spot x-position relative to equally spaced bins (user-defined number of vials) between the minimum and maximum x-values. An optimization plot is created to illustrate the desired ROI and bin assignments, saved with the ‘*ROI*.*png*’ suffix while the data frame containing the filtered spots and their vial assignments contains the ‘*filtered*.*csv’* suffix.

### Calculating climbing velocity, via local linear regression

The mean y-position for all spots in a vial is calculated for each frame. A sliding window is applied to the mean y-position vs. time (velocity) curve to calculate the most linear (greatest regression coefficient) segment of the curve, via local linear regression. The slope of this segment is considered the vial’s velocity (Olito et al., 2017)(Fig. 1E). In videos where the p-value for the regression is not significant (*P* > 0.05), the slope is set to 0.

If conversion factors for pixels-to-cm and frames-to-seconds are supplied and the GUI box is checked or the ‘convert_to_cm_sec’ variable is set to ‘True’, then the plots and final slopes will be converted from pixels per frame to cm per second. This conversion allows for the appropriate comparison across filming setups.

Once all analysis parameters are optimized, the user should save the configuration file (cfg suffix) using the button in the GUI. This file contains all the detection presets and can be used so others can replicate results.

### Automated, high-throughput detection of climbing velocity across many videos

Once the detector is optimized, it can be run from the command line:

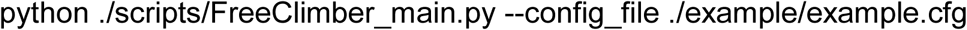

Using the configuration file created in the GUI, the same settings can be applied over all the videos with the specified video type nested in the ‘path_project’ path. The command line interface has several optional arguments for processing subsets of videos (all, unprocessed, or a custom list), generating optimization plots, and preventing the final concatenation of all results files. For more information, please consult the tutorial.

Files containing regression results (including slopes) for each vial in a video are saved with the ‘*slopes*.*csv*’ suffix and are concatenated into a single file after all videos are processed. This file is simply the ‘*results*.*csv*’ file is useful for separate statistical analysis and resides in the ‘path_project’ folder.

### Power Tower: the Drosophila treadmill

The Power Tower automates the process of repeatedly eliciting the negative geotaxis (climbing) startle response, effectively acting as a treadmill (Fig. S3) (Sujkowski et al., 2018; Tinkerhess et al., 2012). Experimental and control flies on the Power Tower were set up in glass vials with food. Flies allowed to “exercise” were placed in vials with the foam stopper at the top to allow climbing, while their “unexercised” control siblings were placed in vials with the foam stopper 1 cm from the food to limit mobility. Flies were knocked down once every 15 seconds while on the Power Tower.

### Longitudinal exercise training program

A longitudinal study over the course of three weeks was conducted with male flies from six mitochondrial haplotypes listed above. Male flies, aged three days post-eclosion, were divided into two groups of 12 vials containing 20 flies under light-CO_2_ anesthesia. Flies were conditioned on weekdays for 2 hours the first week, 2.5 hours the second week, and 3 hours the third week (Piazza et al., 2009), and assayed for climbing performance using the RING assay (Gargano et al., 2005) at the same time each training day before being exercised. Flies were assayed and tested on weekdays and given weekends to recover.

### Endurance exercise fatigue testing

A separate cohort of male flies, aged and collected similarly to the longitudinal cohort, was used to study the mitotypes’ ability to resist endurance climbing fatigue. However, four vials containing 25 flies were set up on the Power Tower (similar to above) and either allowed to exercise (fatigued) or not allowed to exercise (rested). Flies’ initial climbing performances were assayed before being placed on the Power Tower for six consecutive hours and then assayed hourly.

### Statistical analysis on longitudinal data

ANOVA of repeated measures was conducted using the statsmodels (v.0.10.0) module in Python. The ANOVA was used to quantify significant differences between mitochondrial haplotypes, exercise conditions, and the interaction between the two. This test was conducted using the absolute velocities and the normalized climbing index, which represents the climbing velocity normalized by the average velocity from the initial time point.

## Results and discussion

### Local linear regression outperforms a time-based cutoff for climbing velocity

The mean vertical-position vs. time curve is generally concave (Fig. 2A-C) with progress occasionally lagging in the first several frames as flies react to the stimulus, and plateauing at the end as flies reach the top. Traditional manual metrics quantifying the mean vertical position at 2-seconds (or any time point) overestimates the cohorts’ velocity because it assumes flies increase their vertical position linearly. Flies don’t necessarily climb in a straight line, and flies can also have a delayed reaction to the stimulus. This analytical method also assumes flies start at the bottom of the vial. Some flies jump when startled and/or begin at a non-zero starting height, which can create biological noise that is amplified if only a single frame is considered for a time or position-based cutoff. Regardless, reducing a 3D object to a 2D image causes issues as depth is translated into height. Flies starting at the bottom-front of the vial may have a different starting height than those starting at the bottom-back.

**Figure 2.**
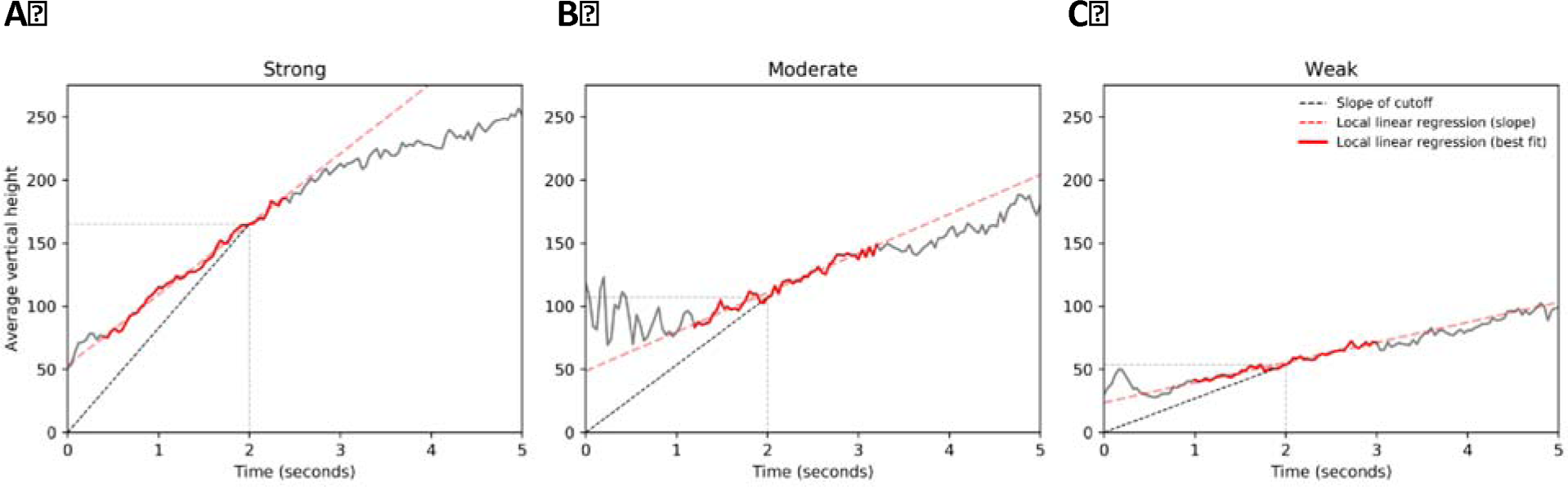
Local linear regression provides a more objective measure of climbing performance. Mean vertical-position vs. time (velocity, solid gray line) plots for a cohort of flies measured at (A) 5, (B) 10, and (C) 20 days post-eclosion. The slope of the standard, time-based cutoff at 2 seconds (black dashed line) has a steeper slope than the line of best fit (red dashed line) during the most linear two seconds of a five second climb (red solid line).

A local linear regression is an appropriate workaround to many of these issues. Here, climbing performance is the slope (velocity) of the most linear portion (greatest regression coefficient over n-consecutive frames) of the mean vertical-position vs. time curve. This approach reduces noise, generates more repeatable and reliable results, and is a more objective measure of performance vs. a cutoff-based method. It can also handle unpredictable climbing behaviors (jumping or falling) and some undetected flies because each fly only has a fractional contribution to overall height. In other words, the local linear regression accounts for context of spots within and across frames, rather than relying on an arbitrary, cutoff-based measure.

### Climbing performance easily quantified for longitudinal studies

Once detection parameters are optimized, FreeClimber can batch process videos. Previous studies demonstrate climbing performance can be affected by genotype (Gargano et al., 2005; Holmbeck et al., 2015; Lavoy et al., 2018), environment (Piazza et al., 2009; Tinkerhess et al., 2012), and genotype x environment effects (Holmbeck et al., 2015; Sujkowski et al., 2018). Accordingly, we tested a set of six, phylogenetically diverse (Ballard, 2000; Montooth et al., 2009), mitochondrial-nuclear (mitonuclear) introgression flies (mitotypes; Fig. 3A). These mitotypes were derived from four different *Drosophila* species: *D. melanogaster* (subtypes: *OregonR* (*OreR*) and *Zimbabwe53* (*Zim*)), *D. simulans* (subtypes: *siI* and *siII* (*sm21*)), *D. mauritiana* (*mall*), and *D. yakuba* (*yak*). All were paired with a common *D. melanogaster* (*w*^*1118*^) nuclear background (mito;nuclear). Four of these lines (*OreR;w*^*1118*^, *siI;w*^*1118*^, *sm21;w*^*1118*^, and *Zim;w*^*1118*^) were previously shown to have weak to moderate climbing performance abilities (Sujkowski et al., 2018), while two (*yak;w*^*1118*^ and *maII;w*^*1118*^) were previously untested.

**Figure 3.**
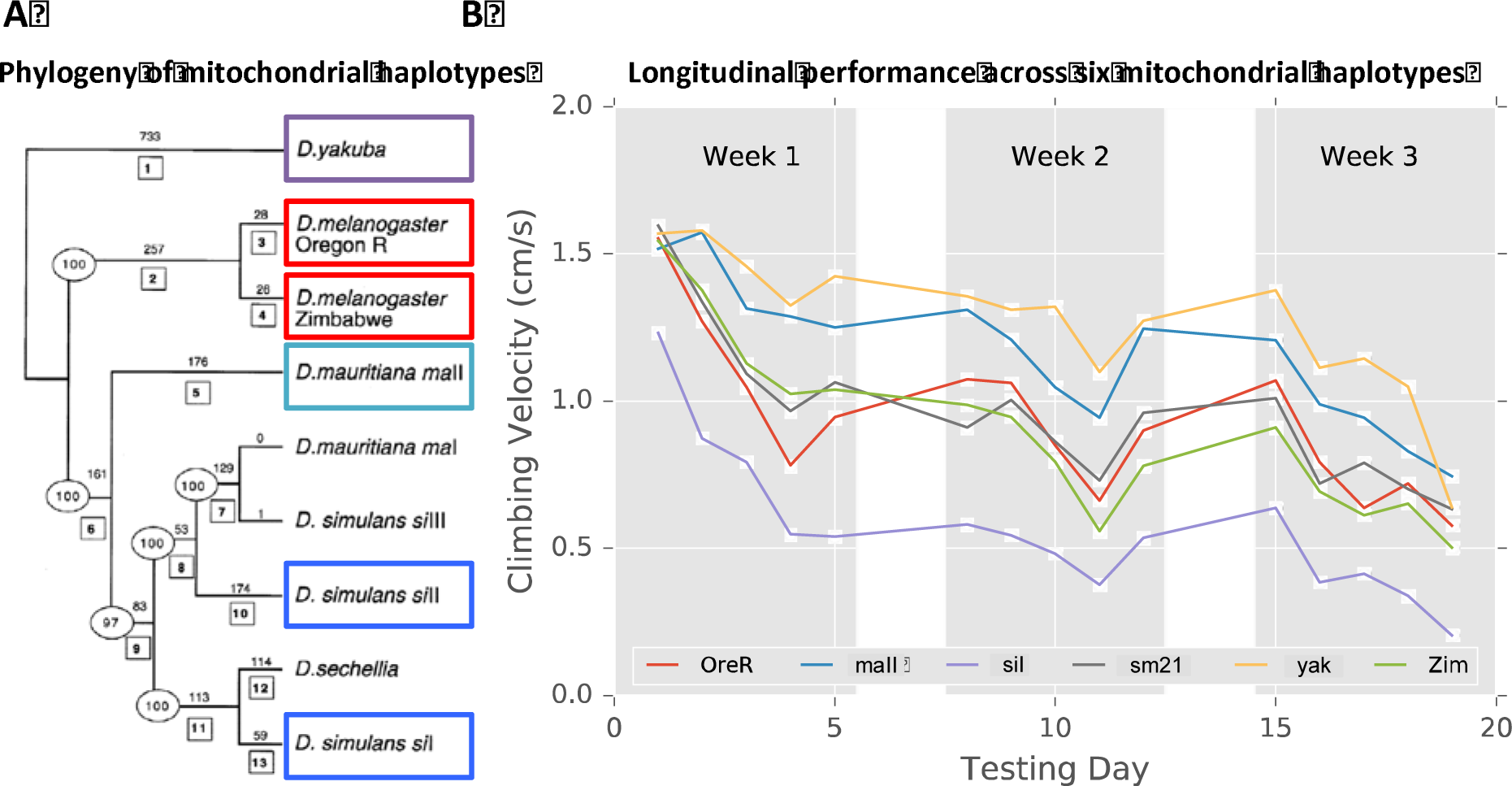
Mitochondrial haplotypes show a differential response to endurance exercise training and resistance to endurance fatigue. (A) Phylogenetically distinct mitochondrial haplotypes were derived from four clades (*D. melanogaster*, red; *D. simulans*, blue; *D. mauritiana*, cyan; and *D. yakuba*, purple). Figure modified from (Ballard, 2000) with *sm21* as a subgroup of *siII*. (B) These mitochondrial haplotypes, on a common nuclear background (*D. melanogaster w*^*1118*^), were subjected to a three-week endurance exercise training program. Flies were tested on weekdays (light gray) before exercise conditioning on a Power Tower, and allowed to rest on weekends (dark gray). Points represent the mean velocity for all flies of a genotype across replicates (n = 1007 videos total). Since there was no significant conditioning effect (*P* = 0.83; Table S2), conditions were averaged together. Most mitotypes began at roughly the same starting velocity, though differed in their age-associated performance decline; *yak;w*^*1118*^ (*yak*, yellow) and *maII;w*^*1118*^ (*mall*, blue) were the strongest overall, while *siI;w*^*1118*^ (*siI*, purple) was the weakest. Note that the color codes differ between panels A and B.

We conducted a longitudinal experiment asking whether mitochondrial haplotypes respond differently to an exercise-conditioning program. We conditioned 12 cohorts (6 mitotypes x 2 training conditions) of 20 male flies following a prescribed training protocol (Sujkowski et al., 2018; Tinkerhess et al., 2012), comparing conditioned cohorts’ daily climbing performance against unconditioned controls. Flies experienced age-associated declines in climbing performance (Fig. 3B) that was significant by mitotype (*P* < 0.0001), but not for conditioning (*P* = 0.83), or mitotype x conditioning effects (*P* = 0.26)(Fig. S3A, Table S2). When normalizing performance by their respective, mean first day performance (normalized climbing index), mitotype (P < 0.0005) and mitotype x conditioning (*P* < 0.0001) were both significant. While there was no significant exercise conditioning effect, the unconditioned flies generally outperformed their conditioned counterparts. This would suggest exercise training is stressful and not always beneficial for the flies. We previously demonstrated mitotypes on the *w*^*1118*^ nuclear background are not sensitive to exercise conditioning effects (Sujkowski et al., 2018), which our results support.

Under the disrupted coadaptation hypothesis (Montooth et al., 2010; Rand et al., 2004), we would expect to see a negative relationship between the divergence between a mito-nuclear pairing and climbing performance. More distantly related pairings have greater opportunity to accumulate mito-nuclear incompatibilities, which would hinder performance. We found *D. melanogaster* pairings (*OreR;w*^*1118*^ and *Zim;w*^*1118*^) were intermediate performers and two divergent lines performed the same (*sm21;w*^*1118*^) or much worse (*siI;w*^*1118*^), supporting the hypothesis. However, the two most divergent pairings (*maII;w*^*1118*^ and *yak;w*^*1118*^) performed the best. A separate study also observed *yak;w*^*1118*^ was longer lived compared to its native mitochondrial-nuclear pairing (*yak;yak*) (Ma and O’Farrell, 2016), providing an independent result supporting our observations and challenging the disrupted coadaptation hypothesis. This finding is surprising, since the *D. melanogaster* and *D. yakuba* sub-species are reproductively incompatible and separated by 7-10 million years of divergence (Consortium, 2007).

Finally, we tested a separate cohort of the same mitotypes’ ability to resist fatigue in a six-hour fatigue assay. We followed a similar Power Tower protocol as the longitudinal study, but instead used four cohorts of 25 flies and had the flies on the Power Tower for one six-hour stretch. We measured initial climbing performance and after each hour. We observed significant mitotype, fatigue, and mitotype x fatigue effects for the absolute velocities, but only a significant fatigue effect after normalizing the slopes to the initial time point (Fig. S3B, Table S2). This fatigue resistance test demonstrates that while the Power Tower may be stressful in a multi-day longitudinal study, it still effectively elicits a consistent climbing phenotype that can slowly fatigue flies over a prolonged period. Interestingly, rested *yak;w*^*1118*^ were strong performers, though their fatigued counterparts had the greatest variation between time points. It is possible that *yak;w*^*1118*^ are strong climbers when undisturbed, but more variable in the climbing performance when stressed.

## Conclusion

FreeClimber is a free and easy-to-use platform for quantifying the climbing velocity for cohorts of flies. It automates the tedious process of detecting and counting flies, and reliably quantifies an objective measure of climbing performance. We applied our platform to measure the longitudinal climbing performance during an exercise-conditioning program and during a resistance to endurance fatigue assay across six mito-nuclear introgression lines. We demonstrate our platform’s ability to identify both strong and subtle differences between genotypes, and its ability to work with longitudinal data sets.

## Acknowledgements

We acknowledge Leann Biancani for her assistance with *Drosophila* husbandry, John Murphy in the Brown University BioMed Machine Shop and Benjamin Wilks for their technical expertise in designing/assembling/maintaining the climbing apparatus, and Kathryn Russo for her time during testing. We thank the Brown University Computational Biology Core (Ashok Ragavendran, August Guang, and Joselynn Wallace) for their technical assistance, and Tom Roberts for providing instruction and materials in the circuit assembly, and for critical feedback on this manuscript.

## Data availability

The source code and example files for FreeClimber, as well as result files used for statistical analysis are available online in a GitHub repository (https://github.com/adamspierer/FreeClimber/tree/JEB_release). The most updated version of FreeClimber can be found at: https://github.com/adamspierer/FreeClimber/.

## List of Symbols and Abbreviations

ecc: eccentricity
GUI: Graphical User Interface
mito-nuclear: Mitochondrial-nuclear
mitotypes: Mitochondrial haplotype
OreR: Oregon R
ROI: Region of Interest
yak: yakuba
Zim: Zimbabwe53

